# Irinotecan induces intestinal atrophy and compromises duodenal motility and mucosal permeability: *in vivo* and *in vitro* evaluations

**DOI:** 10.1101/2025.03.14.643345

**Authors:** Nathalie Arendt, Hans Lennernäs, Femke Heindryckx, Markus Sjöblom

**Affiliations:** Department of Medical Cell Biology, Uppsala University, Uppsala, Sweden; Department of Pharmaceutical Biosciences, Uppsala University, Uppsala, Sweden

**Keywords:** intestinal barrier function, mucosal permeability, motility, chemotherapy, cancer, viability, mucositis, irinotecan, CPT11

## Abstract

**Purpose:** Intestinal toxicity is a significant adverse effect of chemotherapy, with no effective treatments currently available. This study aimed to evaluate the effects of irinotecan (IRT) on; (i) duodenal motility and mucosal barrier function, (ii) histopathological changes along the small intestine and colon, and (iii) faecal water content, to better understand IRT-induced intestinal toxicity, and to support future treatment strategies to minimize these effects.

**Methods:** Sprague Dawley rats were anaesthetized with thiobarbiturate, and a 30-mm long segment of the proximal duodenum with an intact blood supply was perfused *in situ*. The effects of IRT on duodenal motility and blood-to-lumen clearance of ^3^H-mannitol were investigated. In separate experiments, intestinal toxicity was induced in rats by intravenous (i.v.) dosing with IRT. After 72 h post IRT administration, the rats were sacrificed and the small intestine and the colon were excised for histological analysis. Fecal water contents and change in body weight were also measured. In addition, effects of IRT on cell viability were evaluated via Caco-2 cell culture, IRT treatment, and resazurin reduction assays.

**Results:** Intravenous dosing of IRT (60 mg/kg) significantly induced robust and sustained increase in both duodenal motility (+277%) and in mucosal paracellular permeability (+85%). Moreover, IRT in doses of 180 mg/kg and 200 mg/kg (i.p.), induced similar reduction in villus length evaluated 72 h post administration; in duodenum (-52% and -45%, respectively) and jejunum (-37% and -33%), but was without effects in ileum. IRT induced increases in jejunal crypt depth (+48% and +41%), but did not significantly influence the crypt depth in duodenum, ileum or colon. IRT induced increases in fecal water content by +19% and +16%, respectively, and caused significant body weight reductions by 10% and 13%, respectively. IRT exposing of Caco-2 cells resulted in a significant decrease in viability, and a resulting IC_50_ value of 1289 μM.

**Conclusions:** Our novel results show that IRT highly increases duodenal motility and impair mucosal barrier function. In addition, IRT induces diarrhoea, and demonstrates histopathological in the proximal small intestine. These findings contribute to the basic physiological understanding, and provide potentially new innovations for treatment of IRT-induced intestinal toxicity.

## INTRODUCTION

Cancer is one of the leading causes of death worldwide and resulted in almost 10 million cancer-related deaths in 2020, a number estimated to increase to over 16 million by 2040 [1]. Chemotherapy is a cornerstone of cancer treatment, often serving as the first-line therapy for many malignancies [2]. However, while these potent cytotoxic agents effectively target rapidly dividing tumour cells, they also affect proliferating non-tumour cells, particularly epithelial cells in the gastrointestinal (GI) tract and hematopoietic cells, leading to significant side effects [3,4].

Irinotecan (IRT, CPT-11), a semi-synthetic derivative of camptothecin, is a key chemotherapeutic agent used to treat various malignancies, including colorectal, gastric, pancreatic, and ovarian cancers [5]. As a broad-spectrum cytotoxic drug, IRT exerts its antitumor activity primarily by inhibiting topoisomerase I (Topo1), an enzyme involved in DNA replication and transcription [6]. By stabilizing the Topo1-DNA cleavage complex, IRT induces DNA strand breaks, ultimately triggering apoptosis in dividing cells. While Topo1 inhibition is well established as IRT’s primary mechanism, emerging evidence suggests that IRT and its active metabolite may also exert anti-proliferative effects through Topo1-independent pathways [5].

Despite its efficacy, IRT treatment is frequently limited by dose-dependent toxicities, particularly GI complications and neutropenia. Among these, diarrhea is a major dose-limiting side effect, occurring in approximately 40% of patients. Chronic or severe diarrhea may not only necessitate dose reductions, treatment discontinuation and significantly decrease the patient’s quality of life, it can also pose a life-threatening risk [7]. Although IRT remains a cornerstone of cancer therapy, further research into its toxicity mechanisms is needed to optimize its clinical utility while minimizing adverse effects.

The pathophysiology of IRT-induced diarrhoea is complex and multifactorial, involving direct epithelial damage, pro-inflammatory mediator release, and alterations in gut microbiota [8,7]. Conventional antidiarrheal agents provide only limited relief, highlighting the need for a deeper understanding of the underlying mechanisms. Following i.v. administration, IRT is converted into its active metabolite, SN-38, by carboxylesterases, particularly carboxylesterase 2 (hCE2), which is highly expressed in the small intestine, colon, and liver, with the highest expression levels in the epithelium of the duodenal segment [5,9,10]. Despite these highly generated levels, the effects of SN-38 in the duodenal segment are not well known [9]. In the distal small intestine and in the colon, IRT is associated with a reduction in mucin-producing goblet cells alongside increased mucin secretion, likely due to altered mucin gene expression [11]. Additional damage to the enteric nervous system, including a decrease in enteric ganglia within the myenteric plexus, has also been reported [12]. Understanding these mechanisms is essential for developing strategies to mitigate IRT-induced GI toxicity and improve patient outcomes.

Proximal duodenal epithelial cells secrete bicarbonate at high rates to prevent mucosal injury from gastric secretions and, together with bicarbonate from the pancreas, for providing a neutral intraluminal environment for activation of digestive enzymes [13–15]. In addition, an intact regulation of duodenal mucosal paracellular permeability provides a barrier function by preventing harmful agents from entering the body through this route [16]. This permeability is primarily regulated by epithelial apical tight junctions, which can rapidly reorganize in response to various physiological and/or pathophysiological stimuli. The blood-to-lumen clearance of radiolabelled inert probe-markers of various molecular weight, such as ^51^Cr-EDTA or ³H mannitol, is well-established for measuring fast and continuous changes paracellular permeability, i.e. epithelial barrier function [17–23]. In living animals, we have previously shown that duodenal permeability shifts significantly with luminal perfusion of ethanol and acid [24]. Our group also recently showed that enteric intramural excitatory neural reflex involving both nicotinic and muscarinic receptor subtypes mediate an increase in mucosal permeability induced by luminal hypotonicity in rats [17]. Dysregulated tight junction proteins are linked to intestinal mucosal leakage and intestinal mucositis, and may be potentiated by increased intestinal motor activity [16].

The objective of this study is to examine whether administration of IRT induce changes in duodenal functions in the rat, in order to find possible targets for reducing intestinal toxicities. The specific aims were the following: (a) to characterize the effects of i.v. administered IRT on duodenal mucosal permeability and motility; (b) to characterize the dose-related impact of IRT on small and large intestinal epithelial morphology and faecal water content 72 hrs after administration; and (c) to monitor changes in cell viability during IRT exposure.

## MATERIALS AND METHODS

### Ethical statement

All experiments in this study were performed in strict accordance with the Guide for the Care and Use of Laboratory Animals of the National Institutes of Health and were approved by the local ethics committee for experiments with animals (permit 5.8.18-00250/2022) in Uppsala, Sweden. This study follows the ARRIVE guidelines and the study protocol established before the study included information on rat ID and weight, dose(s), dosing time and volumes, sampling times, samples, and sample handling [25]. A humane end-point was defined and animal well-being was monitored twice daily to avoid unnecessary suffering among the included animals. A body weight loss of >20 % was set as a criterion for early termination of the study.

### Chemicals and drugs

The anaesthetic 5-ethyl-5-(1’-methyl-propyl)-2-thiobarbiturate (Inactin®) purchased from Sigma-Aldrich (St. Louis, MO, USA). Sodium chloride (NaCl) and Resazurin sodium salt were purchased from Merck (Darmstadt, Germany). Irinotecan Actavis (solution for infusion 20 mg/ml) and Voltaren® (diclofenac sodium, solution for infusion 25 mg/ml) were purchased from Apoteket AB (Sweden). ^3^H-mannitol was purchased from PerkinElmer Life Sciences (Boston, MA, USA). Ethanol was purchased from Solveco AB (Rosersberg, Sweden). Xylene was purchased from VWR International S. A. S (Rosny-sous-Bois, France). DPX mounting medium, Pertex® was purchased from Histolab® (Askim, Sweden). Hematoxylin mayer and eosin were purchased from Histolab® (Västra Frölund, Sweden). Dulbecco’s Modified Eagle Medium (DMEM), fetal bovine serum (FBS) and penicillin-streptomycin are purchased from ThermoFisher (Stockholm, Sweden).

### Animals

Male Sprague-Dawley rats weighing 250 ± 50 g was obtained from Taconic, Ejby, Denmark, and were delivered to the animal facility at the Biomedical Centre (BMC) at Uppsala University, Sweden. The animals were maintained under standardized temperature and light conditions (12:12-h light-dark cycle; temperature, 21-22°C). The rats were acclimatized in the Animal Department for at least one week before experiments and were kept in enriched caged in groups of two or more with access to tap water and chow (R36, Lantmännen, Kimstad, Sweden) *ad libitum*.

### Experimental and surgical procedure for evaluating acute effects of IRT administration in vivo

Experiments were started by anesthetizing the animal with Inactin® 120 mg·kg^-1^ body weight intraperitoneally. To minimize preoperative stress, anaesthesia was performed within the Animal Department by the person who had previously handled the animals. Then, the rat was immediately transported to the laboratory for surgical procedure.

At the laboratory, the animals were tracheotomized with a flexible tube to facilitate respiration, and the body temperature was maintained at 37.0 ± 0.5°C throughout the experiments using a heating pad controlled by a rectal thermistor probe. The surgical and experimental procedures have been described previously [17,26,27]. A brief summary and some modifications are described here.

A femoral artery and a femoral vein were cannulated with PE-50 polyethylene catheters (Beckton Dickinson and Company, Sparks, MD, USA). The arterial catheter containing 20 IU/mL heparin isotonic saline was used for blood sampling and continuous systemic arterial pressure recordings via connection to a pressure transducer operated PowerLab® system (AD Instruments, Hastings, UK). The vein catheter was used for continuous infusion of ^3^H-mannitol (paracellular mucosal permeability marker) in a 1.5 µCi/mL concentration at a rate of 1.0 mL/h, as well as for drug administration.

A laparotomy was performed, and the common bile duct was catheterized with a PE-10 polyethylene tubing close to its entrance into the duodenum (2-3 mm) to prevent pancreatico-biliary juice from entering the duodenum. Soft silicone tubing (Silastic®, Dow Corning, 1 mm ID) was introduced into the mouth, pushed gently along the oesophagus, guided through the stomach and pylorus, and secured by ligatures 2-5 mm distal to the pylorus. PE-320 tubing was inserted into duodenum approximately 2.5-3.5 cm distal to the pylorus and secured by ligatures. The proximal duodenal tubing was connected to a peristaltic pump (Gilson Minipuls 3, Villiers, Le Bel, France), and the segment was continuously perfused with a 37°C 154 mM sodium chloride solution (saline) at a rate of ∼0.4 mL/min. To complete the surgery, the abdominal cavity was closed with sutures, and the wound was covered with plastic foil.

^3^H-mannitol was administered intravenously as a bolus (3.0 µCi/mL) at this time, and as a continuous infusion of a 1.5 µCi/mL concentration at a rate of 1.0 mL/h. Voltaren i.v. injection 10 mg/kg was given 30 min after completion of the surgery to reverse the surgery-induced paralysis of the intestine [28]. After surgery, 60 min was allowed for the cardiovascular, respiratory, and intestinal function to stabilize before the experiments were commenced.

### Measurement of duodenal motility

Measuring intraluminal pressure was used to assess the duodenal wall contractions. The inlet perfusion tubing was connected, via a T-tube to a pressure transducer, and intraluminal pressure was recorded on a computer. The outlet tubing was positioned at the same level as the inlet tubing. An upward deflection of at least 2 mmHg above baseline was defined as a motor response. Changes in intraluminal pressure were recorded via a digitizer on a computer using PowerLab® and the software Labchart7® (ADInstruments Ldt. Hastings, East Sussex, UK). Duodenal motility was assessed in intervals of 10 min by measure of the total area under the pressure curve (area under the curve; AUC) during the sample period, as described previously [23]. The values given are mean ± SEM of three 10-min intervals (unless stated otherwise).

### Measurement of duodenal mucosal paracellular permeability

After the completion of the surgery, the administered radioactive isotope was permitted one hour for tissue equilibration. The paracellular permeability of the duodenal segment was assessed by the blood-to-lumen clearance of mannitol (a sugar which is non-metabolizable in rats) radiolabelled with ^3^H. Two blood samples (∼0.2 ml each) were collected; the first was collected ten minutes before starting the experiment (at t= -10 min), and the second, after ending the experiment (at t=120). Each blood sample were centrifuged (5000 x g, 5 min) within ten minutes of sample collection, and 50 µL of the resulting plasma was collected for later analysis.

The duodenal segment was perfused with saline at a rate of 0.4 ml·min^-1^, and the perfusate was collected at 10-min intervals. The ^3^H-activity of each blood and perfusate sample was analysed using scintillation (Tri-Carb 2910 TR, Perkin Elmer Life Science, Boston, MA, USA). A linear equation analysis was performed on each sample to obtain a corresponding plasma value for each perfusate sample, to compensate for omission of repeated blood sampling throughout data collection. The blood-to-lumen clearance of ^3^H- mannitol was calculated as described previously [29] (Equation 1), and expressed as (ml·minˉ^1^·100 gˉ^1^).

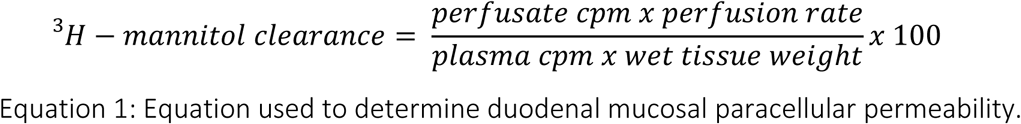

Equation 1: Equation used to determine duodenal mucosal paracellular permeability.

### Experimental protocol

In all of the experiments, the rates of duodenal mucosal paracellular permeability (ml·min^-1^·100 g^-1^), motility (AUC·10 min^-1^), as well as systemic arterial blood pressure (mmHg) were monitored continuously and recorded at 10-min intervals. Control experiments were performed by measuring the parameters above during 120-min perfusions of the duodenal segment with isotonic saline (154 mM NaCl, 300 mOsm·kg^-1^) at a rate of ∼0.4 ml·min^-1^.

In the animals administered irinotecan (IRT), the experiments started with a duodenal perfusion of saline for 30 min to collect basal data. Following this, an intravenous bolus infusion of IRT (60 mg/kg) was administered over a five-minute period. Parameters were collected for another 90 minutes, for a total of 120 minutes of data collection.

### Experimental procedure for evaluating effects of IRT 72h after administration in vivo

#### Study design

This part of the study included three groups of animals. One control group (injected with saline), and two groups that were given a single intraperitoneal (i.p.) injection of either IRT (180 mg/kg) or IRT (200 mg/kg). The rats also received a subcutaneous injection of atropine (0.02 mg) to avoid the immediate and short transient cholinergic side effects of IRT. At 72 h after this administration, the experiments were terminated and the rats were anesthetised (Inactin®, 180 mg/kg). The full colonic content was retrieved, weighed and dried, while duodenal, jejunal, ileal, and colonic tissue samples were taken for morphological evaluation. During deep anaesthesia induced with Inactin® the rats were sacrificed by an intravenous injection of saturated potassium chloride.

The doses of IRT were selected based on previous experience and for clinical translation based on standard doses for IRT [29].

#### *Determining fecal* water content

The faecal water content was measured by extracting the faecal content in colon followed by dehydration of the content. The body weight change was calculated from weight at t=0 and t=72.

#### Histology

Following fixation in 10% neutral-buffered formalin for 24 hours, tissues were longitudinally opened and stored in 70% ethanol. Subsequently, samples underwent dehydration through a graded ethanol series, clearing with xylene, and paraffin infiltration using an automated tissue processor (STP 120 Spin Tissue Processor). Processed tissues were then embedded in paraffin blocks. Paraffin-embedded tissues were sectioned at 5 μm using a rotary microtome (Cool-Cut HM 255 S, Thermo Fisher Scientific, Waltham, MA, USA). Sections were mounted on glass slides (Corning®, NY, USA) and dried overnight on a heated surface.

The tissues were cut into 5 μm sections using a microtome (Microtome Cool-Cut HM 255 S, Thermo Fisher Scientific, Waltham, MA, USA). Tissue sections were then picked up on glass slides (Corning®, NY, USA) and set on a heated surface to set overnight. Hematoxylin and eosin (H&E) staining was performed following standard protocols [30]. Briefly, sections were deparaffinized, rehydrated through graded alcohols, stained with Mayer’s hematoxylin, rinsed, counterstained with eosin, dehydrated, cleared, and coverslipped using DPX mounting medium. After drying overnight, stained sections were examined under a microscope, and images were captured using Zeiss Zen software. Morphometric analysis of villi and crypts was conducted using ImageJ software, measuring structures systematically from left to right in each image to minimize selection bias. Representative images were obtained using a Zeiss Axio Scan.Z1 slidescanner.

### Experimental procedure for evaluating dose-response effects of IRT on cell viability

Caco-2 cells (kindly gifted from Maria Karlgren, Uppsala University) were cultured in Dulbecco’s Modified Eagle Medium (DMEM) supplemented with 10% fetal bovine serum (FBS), and 1% penicillin-streptomycin. Cells were maintained at 37°C in a humidified atmosphere with 5% CO₂. Subculturing was performed at 70-80% confluence using 0.25% trypsin-EDTA, with a seeding density of 2-4 × 10⁴ cells/cm². For the dose-response study, cells were seeded in 96-well plates at a density of 0.1 × 10^5^ cells per well and allowed to adhere overnight. Subsequently, cells were serum-starved for 24 hours, and exposed to varying concentrations of IRT (ranging from 500 to 5000 μM) for 24 hours. Control wells received vehicle treatment only. Cell proliferation was monitored via a resazurin reduction assay. A 1% resazurin sodium salt solution (R7017-1G, Sigma–Aldrich, Darmstadt, Germany) was added in 1/80 dilution to the cells and incubated for 24 h, after which fluorescent signal was measured with a 540/35 excitation filter and a 590/20 emission filter on a Fluostar Omega plate reader.

### Statistical analysis

*In vitro* experiments were done in at least three biological replicates, which we define as parallel measurements of biologically distinct samples taken from independent experiments. Technical replicates we define as loading the same sample multiple times on the final assay. The *in vivo* experiments were done on at least six independent animals. Outliers were kept in the analyses, unless they were suspected to occur due to technical errors, in which case the experiment was repeated. Values are expressed as means ± SEM, with the number of experiments given in parentheses. The statistical significance of the data was tested by repeated measures analysis of variance (ANOVA). To test the differences within a group, a one-factor repeated measures ANOVA was used, followed by a Tukey post-hoc test. Between groups, a two-way repeated measures ANOVA was used, followed by a Bonferroni post-hoc test.

All the statistical analyses were performed using GraphPad Prism 9.4 software (GraphPad Software Inc., San Diego, CA, USA). A *P-*value less than 0.05 was considered significant.

## RESULTS

### Acute effects on IRT intravenous (i.v.) administration in vivo

In control animals perfused with isotonic saline the mean arterial blood pressure (MABP) was 91 ± 2.5 mmHg (not shown). In the irinotecan (IRT) group the MABP was 79 ± 3 mmHg (not shown). It should be noted that administration of IRT (at t=30) induced a 5-10 min transient decrease in MABP (∼10-15 mmHg), a value that stabilised slightly below the previous baseline value.

#### Duodenal mucosal paracellular permeability

In control animals perfused with isotonic saline, the duodenal mucosal paracellular permeability was stable during the 120 min experimental period. At the start of the experiment, permeability was 0.74±0.11 and at the end 0.73±0.07 ml·min^-1^·100 g^-1^; n=6, *P*>0.05; Fig. 1). To investigate the acute effect of IRT on paracellular permeability, an i.v. injection of IRT at a dose of 60 mg/kg was given. IRT induced an immediate and sustained increase in paracellular permeability significant from baseline, from 0.76±0.12 to 1.37±0.24 ml·min^-1^·100 g^-1^; n=6, *P*<0.01; Fig. 1).

**Figure 1.**
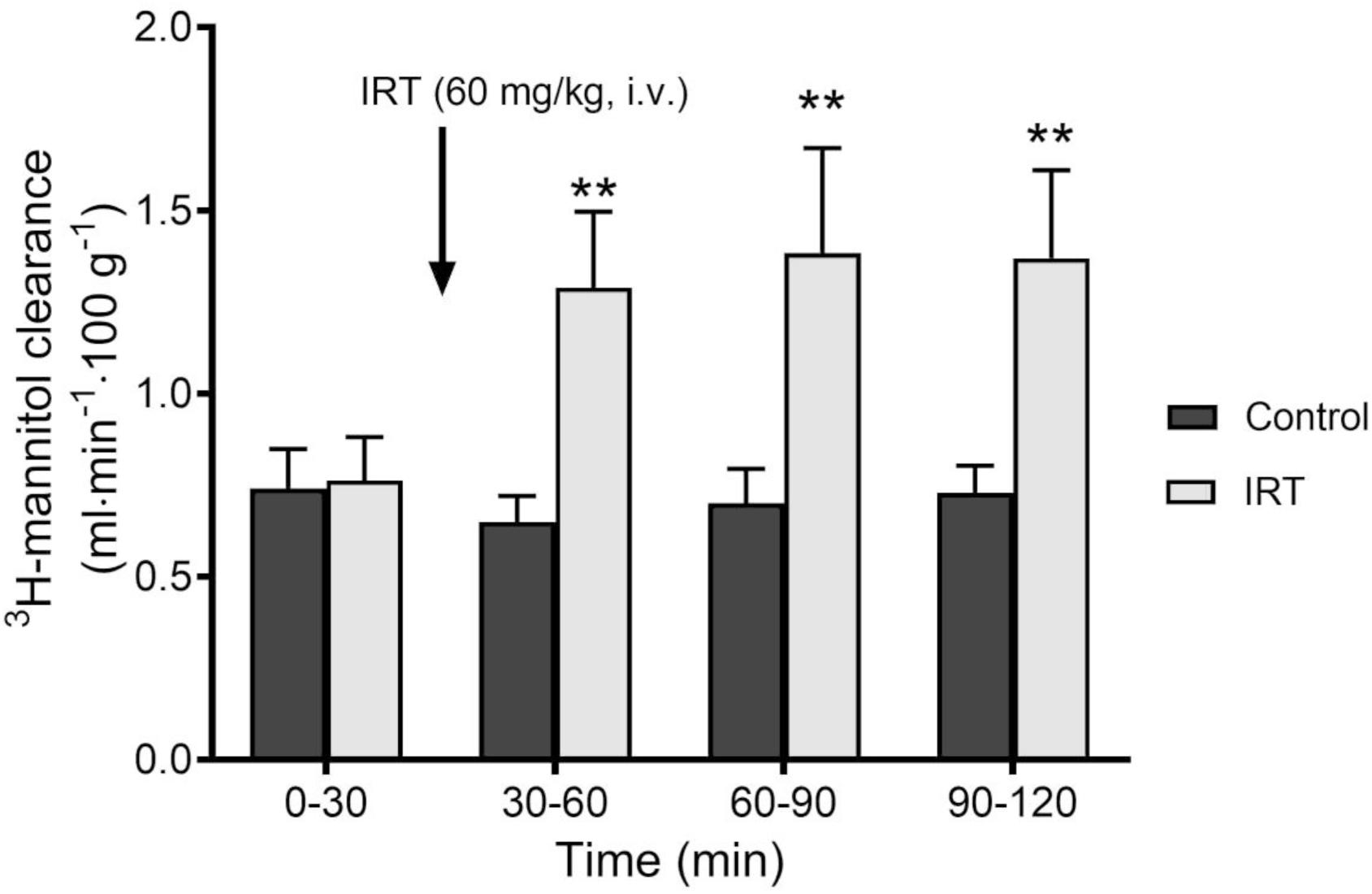
Effects of IRT on duodenal epithelial paracellular permeability. Administration of irinotecan (IRT) (60 mg/kg, i.v.) caused a robust and sustained increase in duodenal epithelial blood-to-lumen clearance of ^3^H-mannitol. In controls the permeability remained stable throughout experiment. Values are mean ± SEM. ** indicates a significant (*P*<0.01) difference compared with the control group.

#### Duodenal motility

In controls, the duodenal motility slightly declined during the 120 min experimental period, from 301 ± 99 to 73 ± 35 AUC·10 min^-1^, values that was not significantly different (n=6, *P*>0.05; Fig. 2). Administration of IRT (60 mg/kg bolus i.v.) induce an instant increase (Fig.3) in duodenal motility from 172 ± 87 to 648 ± 212 AUC·10 min^-1^, and continued to increase over the first 60 min to a maximum value of 733 ± 269 AUC·10 min^-1^ (n = 6, P < 0.01), Fig. 2. During the last 30 min of the experiment motility returned to a value not significantly different from the initial baseline value (208 ± 80 AUC·10 min^-1^ (P > 0.05).

**Figure 2.**
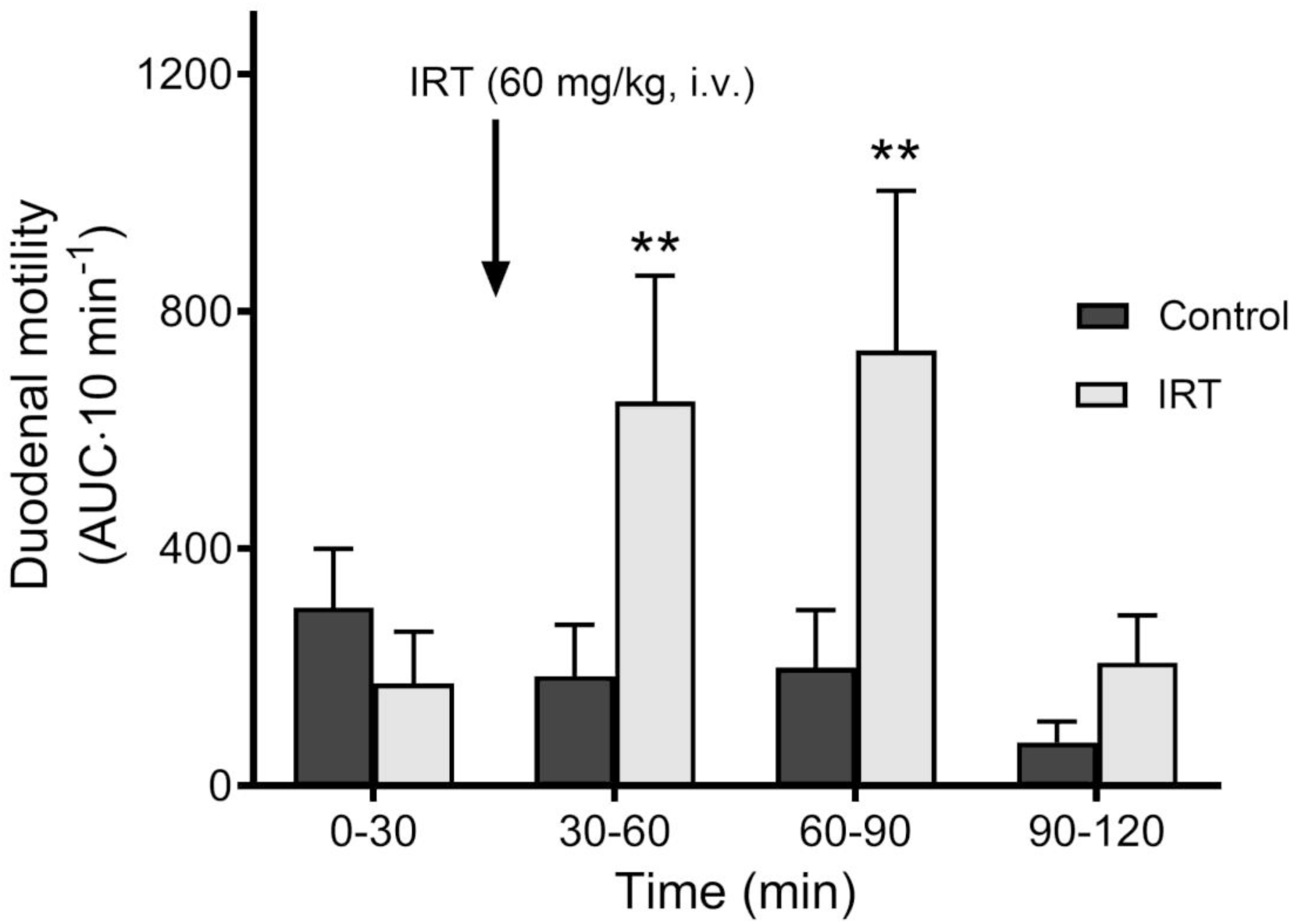
Effects of IRT on duodenal motility. Administration of irinotecan (IRT) (60 mg/kg, i.v.) caused a pronounced transient increase in duodenal motility. 60 min post IRT administration motility returned to the baseline value. In controls the motility stable with a slight decrease throughout experiment. Values are mean ± SEM. ** indicates a significant (*P*<0.01) difference compared with the control group.

**Figure 3.**
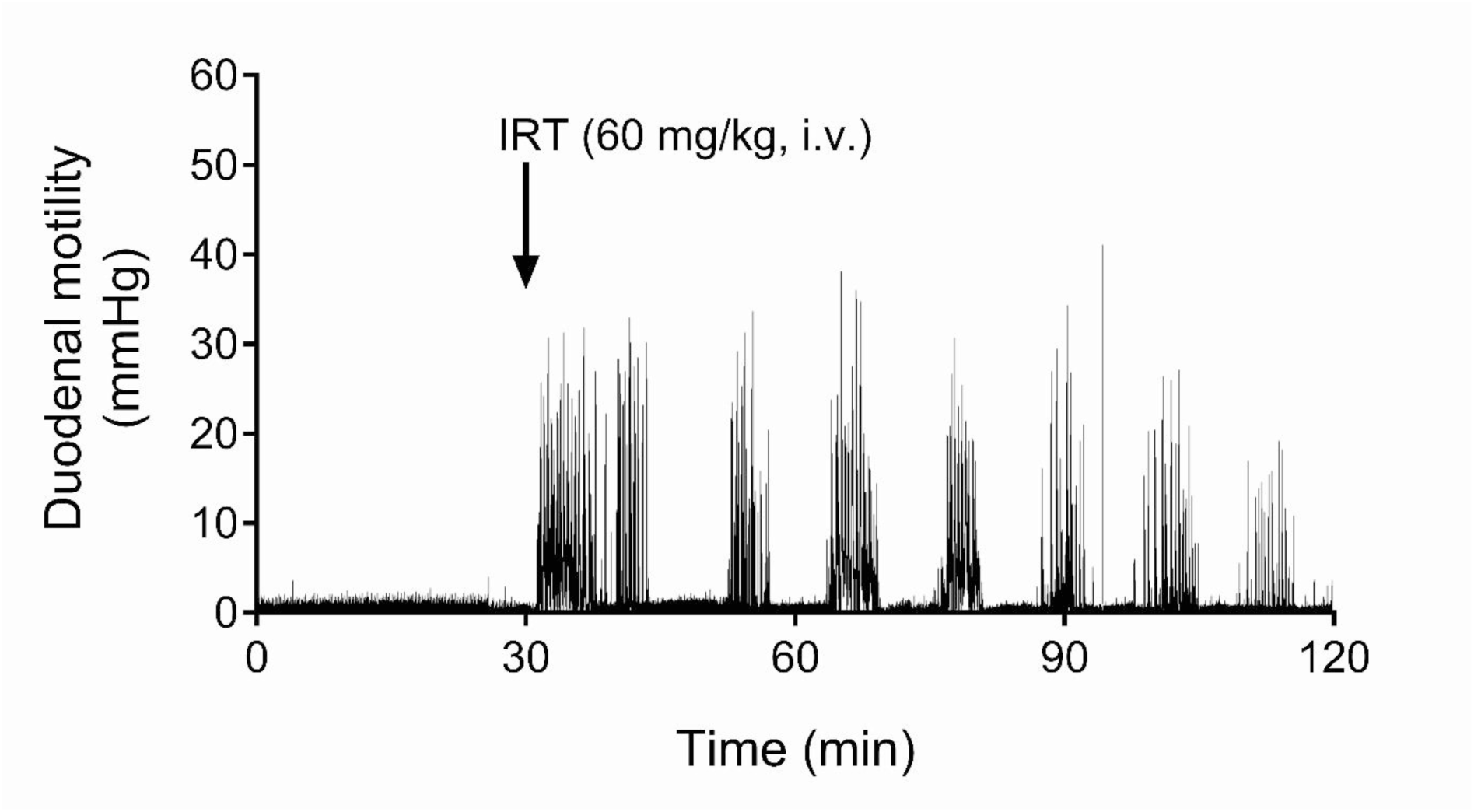
IRT induce an immediate increase in duodenal motility. IRT (60 mg/kg bolus i.v.) induce an increase in duodenal motility within 5 min after administration. The figure illustrates a representative recoding from one experiment.

### Effects of IRT 72h after intraperitoneal (i.p.) administration

#### Histological evaluations

The small intestinal villus height and crypt depth in rats was assessed following a single i.p. injection of a dose of IRT (180 mg/kg, n=6) or IRT (200 mg/kg, n=7). Samples were taken at 72 h post-dosing. The villus height (from villus tip to villus–crypt junction) and crypt depths (defined as invagination depth between adjacent villi) were determined. Histological assessment revealed that a single dose of IRT (180 mg/kg) or IRT (200 mg/kg) significantly reduced the duodenal villus height from 529 ± 23 µm (control, n=8) to 253 ± 16 µm and 289 ± 5 µm, respectively (p < 0.0001), as shown in Fig 4A. Neither IRT (180 mg/kg) nor IRT (200 mg/kg) significantly changed the duodenal crypt depth, Fig. 4B.

**Figure 4.**
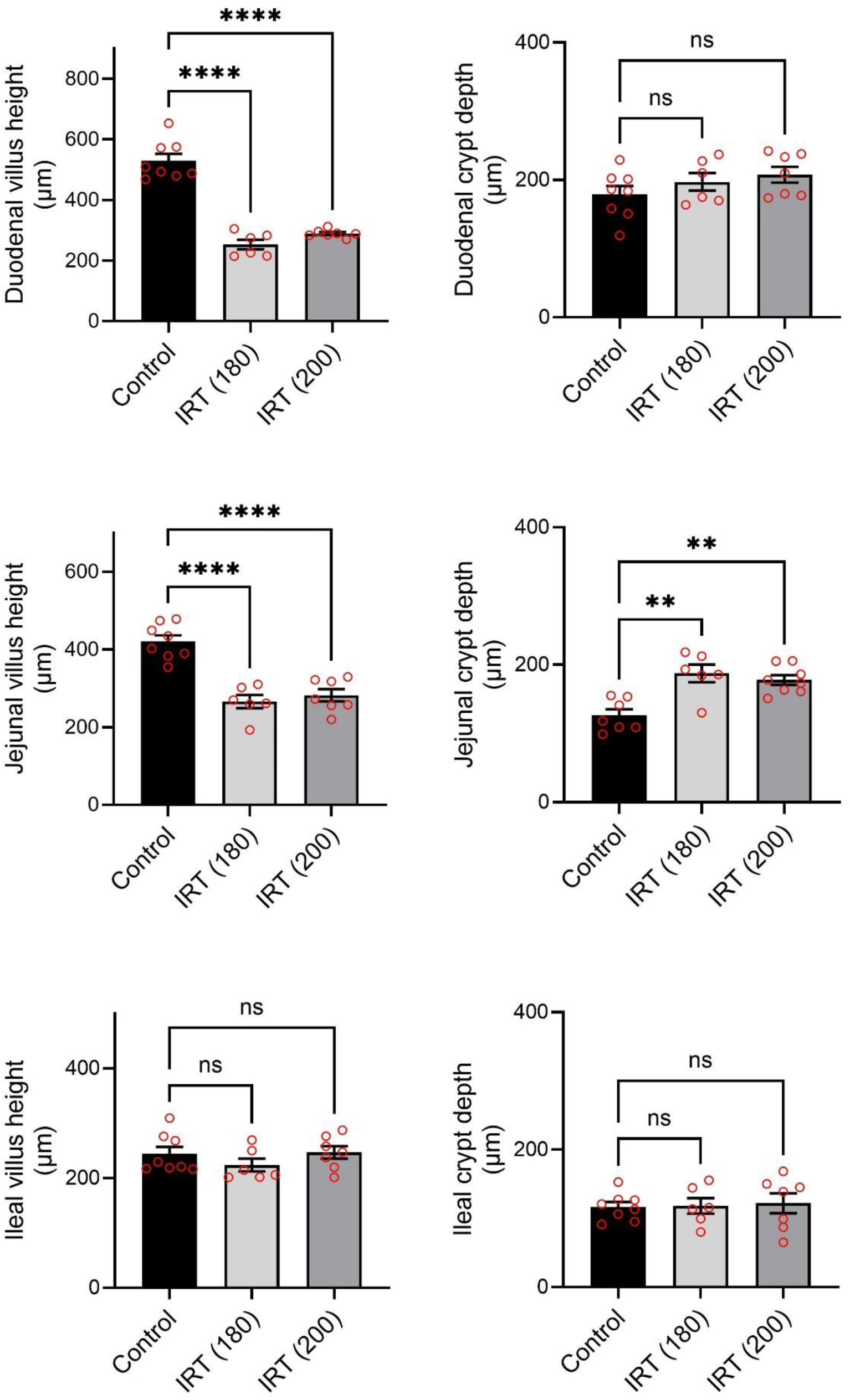
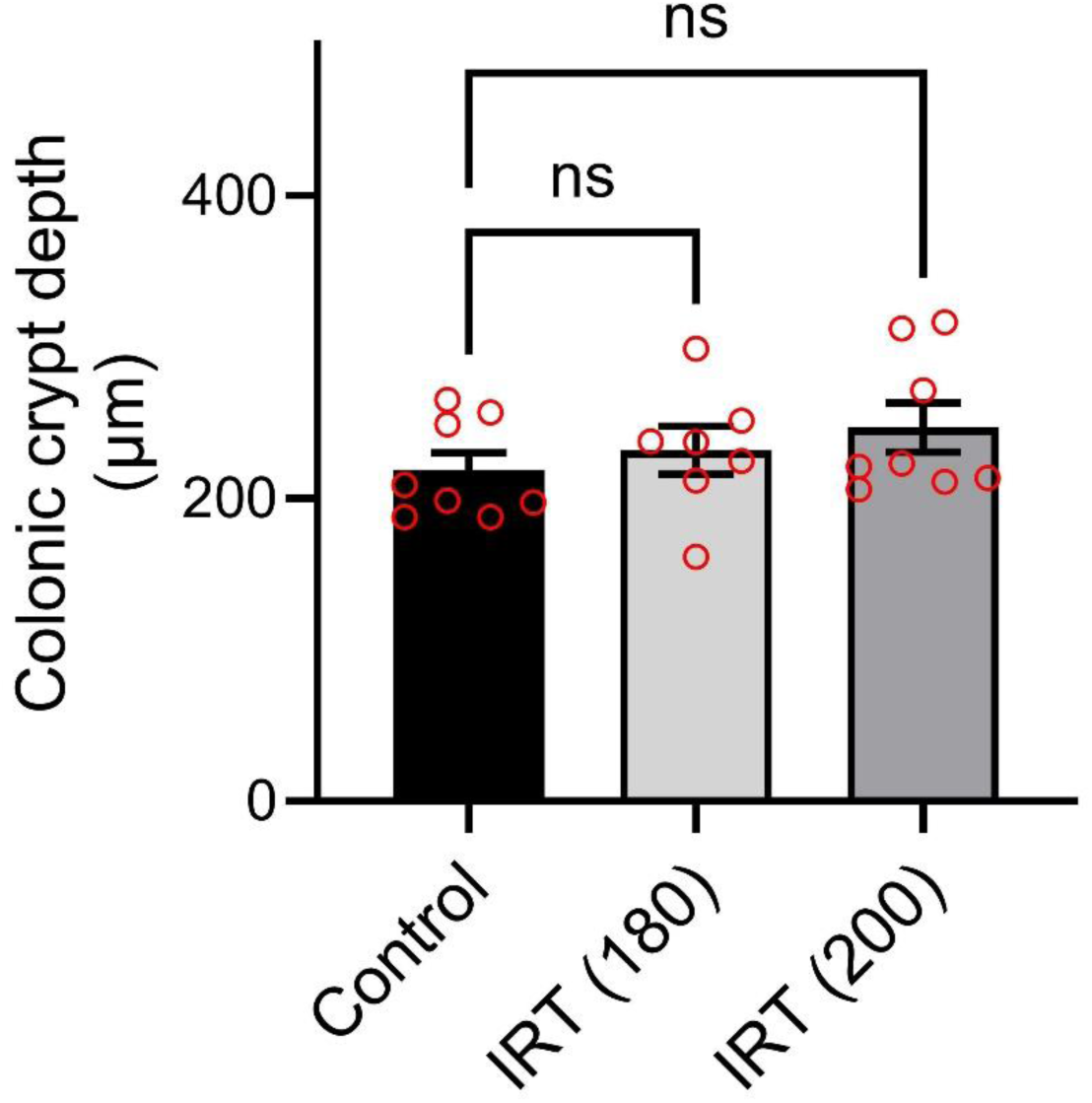
Effects of IRT on intestinal villus length and crypt depth. Effects of irinotecan (180 mg/kg and 200 mg/kg) on a) duodenal villus length, b) duodenal crypt depth, c) jejunal villus length, d) jejunal crypt depth, e) ileal villus length, f) ileal crypt depth, and colonic crypt depth. Values are mean ± SEM. ** and **** indicates significant (*P*<0.01 and *P*<0.0001) difference compared with the control group.

Similarly, IRT (180 mg/kg) and IRT (200 mg/kg) significantly reduced jejunal villus height from 420 ± 16 µm (control, n=8) to 266 ± 17 µm and 282 ± 16 µm, respectively (p < 0.0001), as shown in Fig 4C. IRT (180 mg/kg) and IRT (200 mg/kg) significantly increased jejunal crypt depth, from 126 ± 9 µm (control, n=8) to 187 ± 13 µm and 178 ± 7 µm, respectively (p < 0.01), as shown in Fig 4D. IRT (180 mg/kg) and IRT (200 mg/kg) did not affect ileal villus height (Fig. 4E), ileal crypt depth (Fig. 4F), or colonic crypt depth (Fig. 4G). Representative images are shown for the duodenum (Fig.5A-C), jejunum (Fig.5D-F), ileum (Fig.5G-I), and colon (Fig.5J-L).

**Figure 5.**
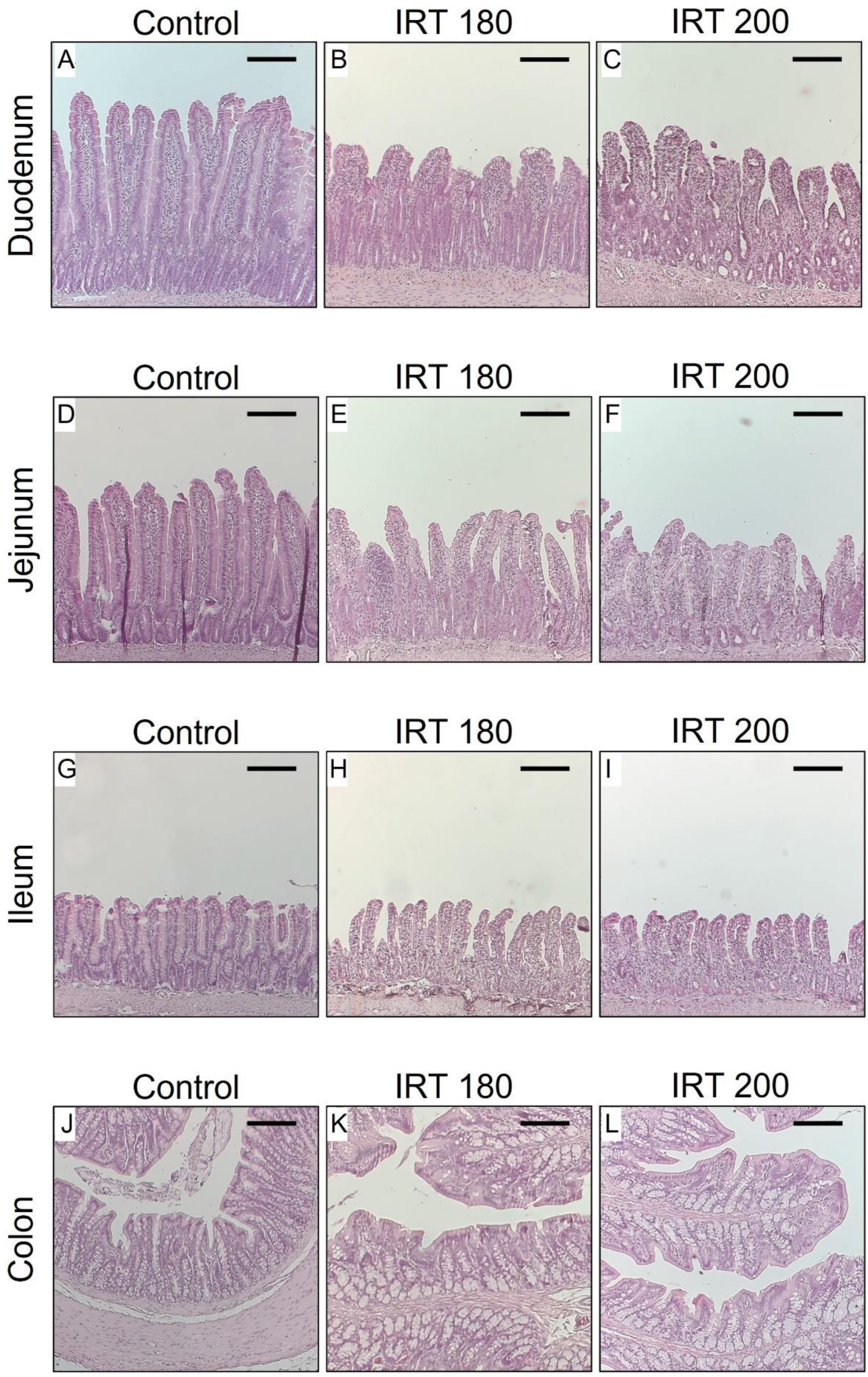
Hematoxylin-eosin staining of the rat intestine. Different specimens from the morphological examination after hematoxylin-eosin staining of the rat intestine 72 h post administration with saline or irinotecan (IRT) in doses of 180 mg/kg or 200 mg/kg. Representative images are shown for the duodenum in (A-C), jejunum in (D-F), ileum in (G-I), and colon in (J-L). The scale indicates 200 µm.

#### Colonic faecal water content

The degree of diarrhea was determined by collecting fecal contents of the colon into plastic tubes. After drying, the water contents could be quantified, calculated and expressed as percent of water. The results at 72 h show that both IRT (180 mg/kg) and IRT (200 mg/kg) increased the fecal water contents from 68 ± 1.1 % (control animals) to 81 ± 1.9 % and 79 ± 3.4 %, respectively (p < 0.01), as shown in Fig. 6A.

**Figure 6.**
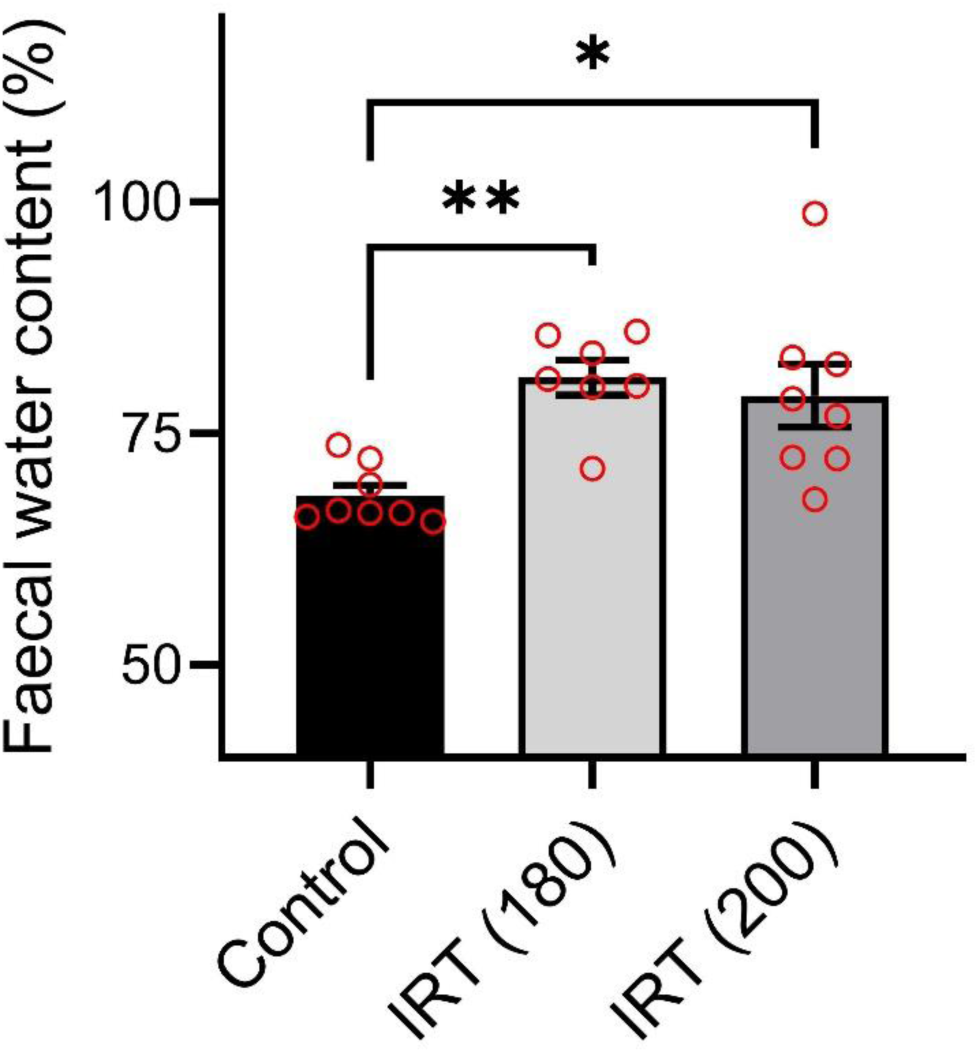
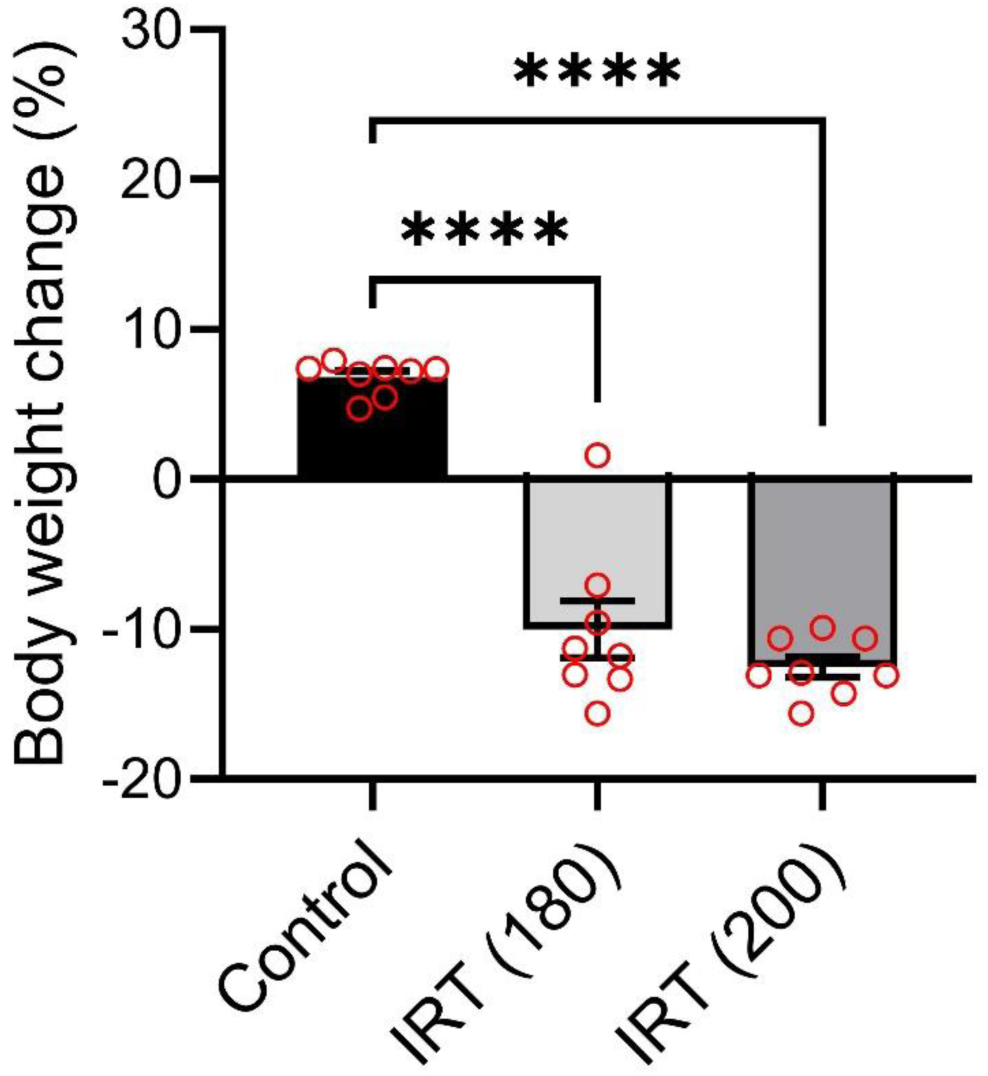
Effects of IRT on faecal water content and body weight. a) Colonic faecal water content in % and, b) change in body weight, 72 h after dosing with irinotecan (IRT) in doses of 180 mg/kg or 200 mg/kg. Values are mean ± SEM. * and ** indicates significant (*P*<0.05 or *P*<0.01, respectively) difference compared with the control group.

#### Changes in body weight

Animals given IRT (180 mg/kg) and IRT (200 mg/kg), demonstrated a significant reduction in body weight 72 h after administration, -10.1 ± 1.9 % and -12.5 ± 0.7 % respectively, compared to controls (+6.8 ± 0.4 %), Fig. 6B.

### In vitro – dose-response effects of IRT on cell viability

CaCo-2 cells were exposed to IRT at concentrations ranging from 500 to 5000 μM, and cell viability was assessed using the alamarBlue assay. The effect of IRT on cell viability was concentration-dependent, resulting in a calculated IC₅₀ value of 1.2 mM (Fig. 7). These findings confirm that IRT exerts a dose-dependent cytotoxic effect on intestinal epithelial cells *in vitro*.

**Figure 7.**
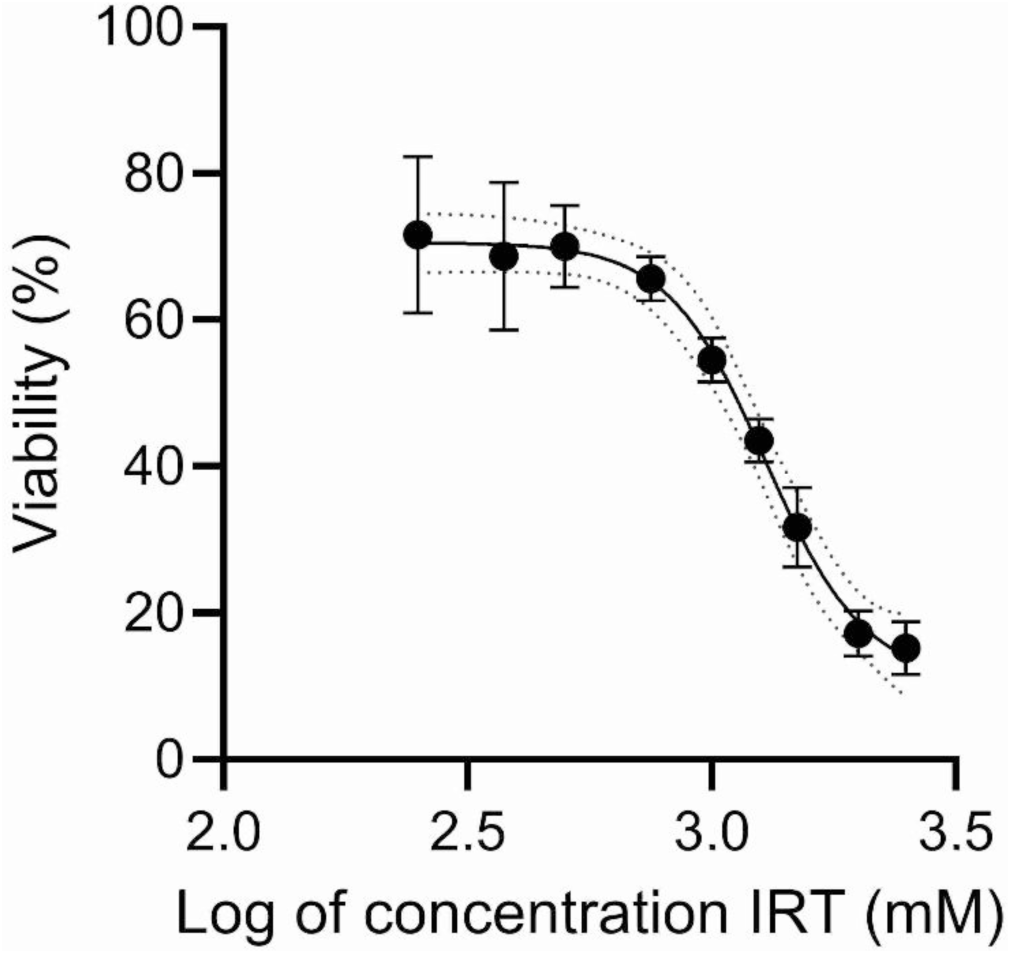
Effects of IRT on CaCo-2 cell viability. Graph representing % cell viability versus logarithm of the concentration of IRT. IC50 was calculated to 1.3 mM in CaCo-2 cells.

## DISCUSSION

Chemotherapy-induced intestinal toxicity is a serious side effect that leads to significant disruptions in gut function. It is marked by increased intestinal permeability, inflammation of the mucosa, digestive impairments, fluid imbalance, and difficulties in absorbing electrolytes and nutrients [3,4]. Additionally, it can cause microbial imbalances (dysbiosis) and changes in intestinal motility, resulting in a range of GI symptoms and broader systemic health complications [31,4,32]. Despite extensive research using animal models, only a limited number of findings have successfully translated into clinical strategies for managing chemotherapy-induced intestinal toxicity [33,34]. However, the fundamental insights gained from these pre-clinical models play a crucial role in shaping our current understanding of the pathophysiology, prevention, and treatment of intestinal toxicity.

For the first time, we demonstrate that IRT induces distinct and interconnected effects on intestinal physiology, linking its acute impact on duodenal motility and mucosal barrier function with delayed structural damage and increased faecal water content. Firstly, IRT administration in rats leads to significant increases in duodenal mucosal paracellular permeability and motility, which happened shortly after intravenous infusion. Additionally, 72 hours post-administration IRT induces villus atrophy predominantly affecting the proximal small intestine. This is an interesting observation, as this can be correlated to that the duodenal epithelium has the highest enzymatic capacity to metabolize the prodrug IRT to the active substance SN-38 [9,10]. Furthermore, we found that IRT treatment results in increased faecal water content, consistent with previous studies [35–37,7]. Altogether, these observations collectively highlight the multifaceted impact of IRT on gut function, providing new insights how the mechanisms and their timing contribute to chemotherapy-induced intestinal toxicity.

Intestinal paracellular permeability is largely regulated by tight junctions, which form a dynamic, selective barrier between epithelial cells within the apical junctional complex. The major rate-limiting pathway for passive transport of solutes with a low molecular mass and fluid across the intestinal epithelium is the paracellular route, which is regulated by tight junctions within the apical junctional complex [38,19]. Tight junctions consist of a complex protein structure that forms dynamic, selective pores in the epithelium that control permeability [16]. To our knowledge, no continuous measurement has been reported of *in vivo* duodenal permeability of low-molecular-mass probes following administration of IRT. These measurements can contribute to fill a knowledge-gap on how IRT mediates mucosal injury and barrier dysfunction. In this study, we measured duodenal mucosal paracellular permeability, by continuously monitoring the blood-to-lumen clearance of ^3^H-mannitol, a well characterized probe that gives a valid estimate on changes in epithelial paracellular permeability. Smaller probes, such as mannitol, are likely to be transported paracellularly along the whole crypt-villus axis [18]. After administration of IRT, an increase in duodenal permeability was observed within 5-15 minutes, suggesting that IRT causes dilation of the epithelial paracellular pathway. The duration of this increase in permeability remained high during the rest of the experiment. The most rational explanation for the almost instant increase in permeability, or leakage, is that IRT directly, or indirectly, affects the intracellular signaling pathways in the intestinal enterocytes, causing changes in the intercellular adhesion between cells. The sustained increase in permeability is likely due to epithelial damage, as previously reported toxic effects of IRT on intestinal epithelial cells have been associated with dysregulation of tight junctions. Supporting this, we observed a concentration-dependent cytotoxic effect of IRT on CaCo-2 cells, with a significant reduction in cell viability at increasing doses. Interestingly, previous studies on irinotecan’s cytotoxic effects on CaCo-2 cells have shown variability in cell survival outcomes based on concentration and exposure duration. A study by Kaku et al. (2015) reported that irinotecan does not induce apoptosis in CaCo-2 cells, but their experiments were conducted with a maximum concentration of 30 µM over 24 and 48 hours, which is significantly lower than the concentrations used in our study [39]. Another *in vitro* cell study observed a 40% reduction in cell survival at approximately 170 µM and complete cell death at 1477 µM after 24 and 48 hours [40]. Overall, this suggests that irinotecan’s cytotoxic effects are highly dose-dependent, with significant variations in cell survival at different concentrations and likely also influenced by initial seeding densities and drug exposure times [40]. This loss of viability in intestinal epithelial cells after *in vitro* exposure of IRT to a colon carcinoma cell line may explain the observed increase in duodenal permeability in the rat model, as cell death and structural disruption likely contribute to barrier dysfunction. Together, these findings reinforce the notion that IRT not only induces transient functional changes in epithelial permeability, but also causes direct epithelial injury, which may play a critical role in chemotherapy-induced mucosal toxicity.

Further supporting this, our previous studies have shown that paracellular permeability returns to baseline level after cessation of the duodenal perfusion with the potent barrier breaker ethanol (10-15%), indicating that the response on permeability is reversible when the toxicity does not induce damage [24]. Similar dynamics has been shown *in vitro* using Caco-2 cell line, that noncytotoxic concentrations of ethanol increases paracellular permeability by displacement and disruption of tight junction proteins and that the paracellular gaps re-closed when ethanol was removed [41]. We have previously found that a single dose of IRT does not have any impact on jejunal mucosal paracellular permeability at a time-point of 72 h after administration, although the dose given was almost three times higher than in the present study (150 mg/kg vs 60 mg/kg) [29]. This suggests that in healthy rodents, IRT-induced increases in small intestinal permeability are only present a fairly short time after administration of a single dose. However, it remains to be investigated how repeated injections of IRT with recovery intervals would affect the permeability, and how experimental cancer models — where both the presence of tumours and the effects of treatment may influence adverse outcomes—would alter these responses. It is also necessary to determine systemic and local intestinal tissue exposure of unchanged prodrug IRT and active drug SN-38 that is locally formed in order to be able to establish a concentration-effect relationship for these effects for both prodrug and active drug. It is also unclear whether this rather short and transient increase in permeability is sufficient to initiate a sustained and self-perpetuating cycle of mucositis.

Chemotherapy may influence gastrointestinal motility either directly, through effects on smooth muscle cells, or indirectly, by altering the enteric nervous system. However, this remains an underexplored area of research [34]. While intestinal transit time can be obtained from animal models and patients, it provides only limited information on the motility pattern. In contrast, experiments on anesthetized animals with an isolated intestinal segment, preserving both an intact blood supply and enteric neural network - as used in the present study – offer an attractive experimental model where numeral functional mechanisms can be studied simultaneously. An interesting finding in our study was that the duodenal motility was greatly affected by administration of IRT. The increase in motility had an almost immediate (within 5 minutes) onset after injection, an observation that has not been described previously. This effect may, at least in part, be the reason why some patients experience diarrhoea within 30 minutes after the administration. Supporting this, previous studies have shown that IRT can induce early-onset diarrhoea as a consequence of its inhibition of acetylcholinesterase, to induce an acute cholinergic syndrome [7]. The increased levels of acetylcholine, may induce intestinal motor dysfunction due to stimulation of cholinergic muscarine receptors [5], which could explain the rapid increase in motility. However, conversely, we recently showed that administration of bethanechol, a specific muscarinic agonist without any effect on nicotinic receptors, did not affect the duodenal motility in the same experimental model [17]. These findings highlight the complexity of IRT-induced motility changes and warrant further investigation across different intestinal segments. While our study demonstrates increased intestinal motility following IRT administration, Boeing et al. have reported the opposite effect, showing reduced digestive motility in mice treated with IRT [42]. This apparent discrepancy underscores the need for further research to clarify the mechanisms underlying chemotherapy-induced alterations in intestinal motility.

In the present study classical signs of intestinal toxicities were observed 72 hours post- IRT administration, including body weight loss, increased fecal water content, and histopathological changes of the small intestine and colon. IRT induced a marked shortening the duodenal and jejunal villi, while no differences in ileal villi length were observed. A particularly interesting finding was the presence of jejunal crypt hyperplasia in rats sacrificed 72 h after IRT administration. This hyperplasia was not seen in any other segments of the small intestine. A similar hyperplasia was previously reported by Gibson et al 2003 [43], and has been confirmed in rodents by others [44,45], including our group ([36]). The most plausible explanation for this phenomenon is that the jejunum exhibits the fastest increase in mitotic cell numbers, marking the initiation of the recovery process following IRT-induced damage. We found that morphological changes in the colonic epithelium were minimal. However, this does not necessarily mean that the function of the epithelium in the crypts in this segment are not affected by IRT. Previous studies have shown an increase of colonic crypt 24h after administration of doxorubicin (Dekaney et al., 2009).

Chemotherapy-induced diarrhoea can be a severe condition, potentially limiting treatment or posing serious health risks [4,7]. This study’s findings align with previous research, demonstrating that IRT leads to a significant rise in faecal water content—an established marker of diarrhoea in animal studies. Additionally, the rapid and substantial body weight loss observed within 24 hours suggests that the decrease is primarily due to increased faecal output and reduced food and water intake, rather than a direct loss of fat or muscle mass. However, this remains speculative, as we did not measure these factors directly.

The increase in faecal water content during the first 24 h after administration is likely due to a combination of increased small intestinal motility and increased mucosal permeability. However, we assert that the diarrhoea is probably driven by different mechanisms. One proposed mechanism, established in animal models, involves the metabolism of SN-38. When SN-38 undergoes hepatic glucuronidation, it is converted into SN-38 glucuronide (SN-38G) and excreted into the intestinal tract via bile. In the colon, bacterial β-glucuronidases can hydrolyse SN-38G, reactivating SN-38 that causes damage to the epithelium which is believed to contribute to delayed-onset diarrhoea [46].

In conclusion, this study shows that IRT has a significant impact on the regulation of duodenal motility and mucosal barrier function. The observed increase in motility and mucosal permeability may contribute to both the acute and delayed diarrhea associated with IRT treatment. The most pronounced histopathological changes were observed in the proximal small intestine, further reinforcing the idea that early disruptions in motility and permeability may drive IRT-induced intestinal toxicity. Despite its well-documented challenges, including intestinal toxicity, IRT remains a cornerstone of cancer therapy. Therefore, further research is needed to elucidate the relationship between epithelial fluid handling, electrolyte transport, mucosal permeability, and motility in the proximal small intestine. Given that the duodenum and jejunum play a crucial role in the activation of IRT to its active metabolite, SN-38, expanding our understanding of its localized effects is essential. Future studies focusing on these mechanisms could provide valuable insights for optimizing clinical efficacy while minimizing adverse effects, ultimately improving patient outcomes.

## ADDITIONAL INFORMATION

Competing interests None declared.

### Author contributions

N.A. and M.S. conceived and designed the research, prepared figures and drafted the manuscript. N.A. performed experiments. N.A. and M.S. analysed data. N.A., F.H., and M.S. interpreted results of experiments. All authors edited and revised the manuscript. All authors approved the final version of the manuscript and agree to be accountable for all aspects of the work in ensuring that questions related to the accuracy or integrity of any part of the work are appropriately investigated and resolved. All persons designated as authors qualify for authorship, and all those who qualify for authorship are listed.

## Acknowledgement

We thank Dr. Karsten Peters, Josefin Marklund, Saarah Trädgårdh and Clara Andersson for excellent technical assistance.

